# Comparative analysis of metabolic models of microbial communities reconstructed from automated tools and consensus approaches

**DOI:** 10.1101/2023.09.13.557568

**Authors:** Yunli Eric Hsieh, Kshitij Tandon, Heroen Verbruggen, Zoran Nikoloski

## Abstract

Genome-scale metabolic models (GEMs) of microbial communities offer valuable insights into the functional capabilities of their members and facilitate the exploration of microbial interactions. These models are generated using different automated reconstruction tools, each relying on different biochemical databases that may affect the conclusions drawn from the *in silico* analysis. One way to address this problem is to employ a consensus reconstruction method that combines the outcomes of different reconstruction tools. Here, we conducted a comparative analysis of community models reconstructed from three automated tools, i.e. CarveMe, gapseq, and KBase, alongside a consensus approach, utilizing data from two marine bacterial communities. Our analysis revealed that these reconstruction approaches, while based on the same genomes, resulted in GEMs with varying numbers of genes and reactions as well as metabolic functionalities, attributed to the different databases employed. Further, our results indicated that the set of exchanged metabolites was more influenced by the reconstruction approach rather than the specific bacterial community investigated. This observation suggests a potential bias in predicting metabolite interactions using community GEMs. We also showed that consensus models encompassed a larger number of reactions and metabolites while concurrently reducing the presence of dead-end metabolites. Therefore, the usage of consensus models allows making full and unbiased use from aggregating genes from the different reconstructions in assessing the functional potential of metabolic communities.

**Importance:** Our study contributes significantly to the field of microbial community modeling through a comprehensive comparison of genome-scale metabolic models (GEMs) generated via various automated tools, including: CarveMe, gapseq, KBase, and a consensus approach. We revealed substantial structural disparities in model outcomes, primarily attributed to variations in the employed databases. A key finding underscored the substantial impact of the reconstruction approach on the set of exchanged metabolites, emphasizing the necessity for enhanced data integration strategies. The consensus models emerge as a powerful solution, exhibiting improved functional capabilities by encompassing a greater number of reactions, metabolites, and genes. This not only offers a more comprehensive representation of metabolic networks within bacterial communities but also shows promise in reducing variability for more accurate predictions of exchange metabolites. Overall, our research provides a critical framework for refining microbial community simulations, impacting fields from ecology to synthetic biology.

## Intruduction

Microbe-microbe interactions play a crucial role in maintaining microbial diversity, influence metabolic phenotypes, and shape community functionality (1, 2). Therefore, microbial communities and interactions are increasingly studied in agriculture (3), synthetic biology (4), pathology (5), and ecology (6). Microbial interactions are in part achieved by the exchange of metabolites, and they are particularly challenging to study in wild communities (7). As a complementary tool, genome-scale metabolic models (GEMs) provide means to identify and dissect the effect of these interactions.

Constraint-based modeling using GEMs has been used to investigate the activity of different reactions in a metabolic network, including exchange reactions that model interactions between microbes. Numerous studies have employed GEMs to investigate metabolic interactions and functionality within microbial communities, including those found in the human gut (8), termite gut (9), mangrove sediments (10), soil microbial communities (11), and plant root (12). Community-scale metabolic models are typically constructed using: (i) the mixed-bag approach, which involves integrating all metabolic pathways and transport reactions into a single model with one cytosolic and one extracellular compartment; (ii) compartmentalization, where multiple GEMs are combined into a single stoichiometric matrix, with each species assigned to a distinct compartment; (iii) costless secretion, wherein models are simulated using a dynamically and iteratively updated medium based on exchange reactions and metabolites within the community (13, 14). The choice of approach depends on the specific objectives and scenarios. The mixed-bag approach is suitable for analyzing interactions between communities, while the other approaches are more appropriate for understanding interactions between organisms within a community (15).

Regardless of the approach used, *in silico* analysis of metabolism of individual organisms in a community requires access to reconstructed GEMs for all species in the community. Several automated approaches are available for GEM reconstruction, including: CarveMe (16), gapseq (17), and KBase (18). Mendoza et al. (19) conducted a systematic evaluation of reconstruction tools, revealing that each tool offers distinct features. For example, CarveMe enables fast model generation due to their ready-to-use metabolic networks, while gapseq incorporates comprehensive biochemical information by employing various data sources during reconstruction. However, selecting different tools can lead to the construction of alternative networks, introducing uncertainty in the predictions resulting from the constraint-based modeling with these GEMs (20). This uncertainty could be caused by gene annotation, gene-reaction mapping, biomass composition, and environment specification. The structure of the reconstructed network is significantly influenced by the choice of the database of biochemical reactions, and this variation is potentially caused by mis-annotations (21) and hypothetical sequences of unknown function (22). During reconstruction, the inclusion of specific reactions in the model depends on the genomic evidence and the network context, often omitting certain reactions based on the modeling objectives. Furthermore, the use of different namespaces for metabolites and reactions from various data sources can pose challenges when combining GEMs (11, 23), leading to further difficulties in predicting metabolic phenotypes of microbial communities.

Consensus models, formed by integrating different reconstructed models of single species from various tools, have the potential to reduce the uncertainty existed in a single model (24, 25) and can be used to estimate interactions in a community (11). However, a systematic comparison between consensus models and original models in terms of model structure (i.e. the number of reactions and metabolites), the inclusion of genes in the model, model functionality, and the potential exchange of metabolites at the community scale is currently lacking. Here, we conducted a comprehensive analysis of these features for models reconstructed using three automated tools, namely: CarveMe, gapseq, and KBase, and a recently proposed consensus reconstruction proposed (11). Our findings shed light on the advantages and limitations of each approach. Specifically, the analysis revealed that consensus models retain the majority of unique reactions and metabolites from the original models, while reducing the presence of dead-end metabolites. Furthermore, consensus models incorporate a greater number of genes, indicating stronger genomic evidence support for the reactions. These characteristics of consensus models demonstrate their enhanced functional capability and capacity for more comprehensive metabolic network models in a community context.

## Results

### Structural differences in genome-scale metabolic models from two bacterial communities

We utilized a collection of 105 high-quality metagenome-assembled genomes (MAGs) derived from coral-associated and seawater bacterial communities described in Robbins et al. (26) to construct genome-scale metabolic models. GEM reconstruction used three automated approaches: CarveMe. (16), gapseq (17) and KBase (18), to generate draft models. Draft models originating from the same MAG were merged to construct draft consensus models by using a recently proposed pipeline (11). Gap-filling of the draft community models was performed using COMMIT (11) (see Methods).

To compare the structural characteristics of the community models, we examined the number of reactions, metabolites, dead-end metabolites, and genes in the resulting reconstructions (Fig. 1). Genes serve as the fundamental components of GEMs. Inclusion of a gene in the model indicates its association with at least one biochemical reaction, thus affecting the set of metabolites in the models. Our analysis revealed that CarveMe models exhibited the highest number of genes, followed by KBase and gapseq in models of both coral-associated bacterial and seawater bacterial communities. Additionally, gapseq models encompassed more reactions and metabolites compared to CarveMe and KBase models, potentially indicating that many genes in gapseq models are associated with multiple reactions. However, gapseq models also exhibited a larger number of dead-end metabolites, which may affect on the functional characteristics of the models.

**Fig 1.**
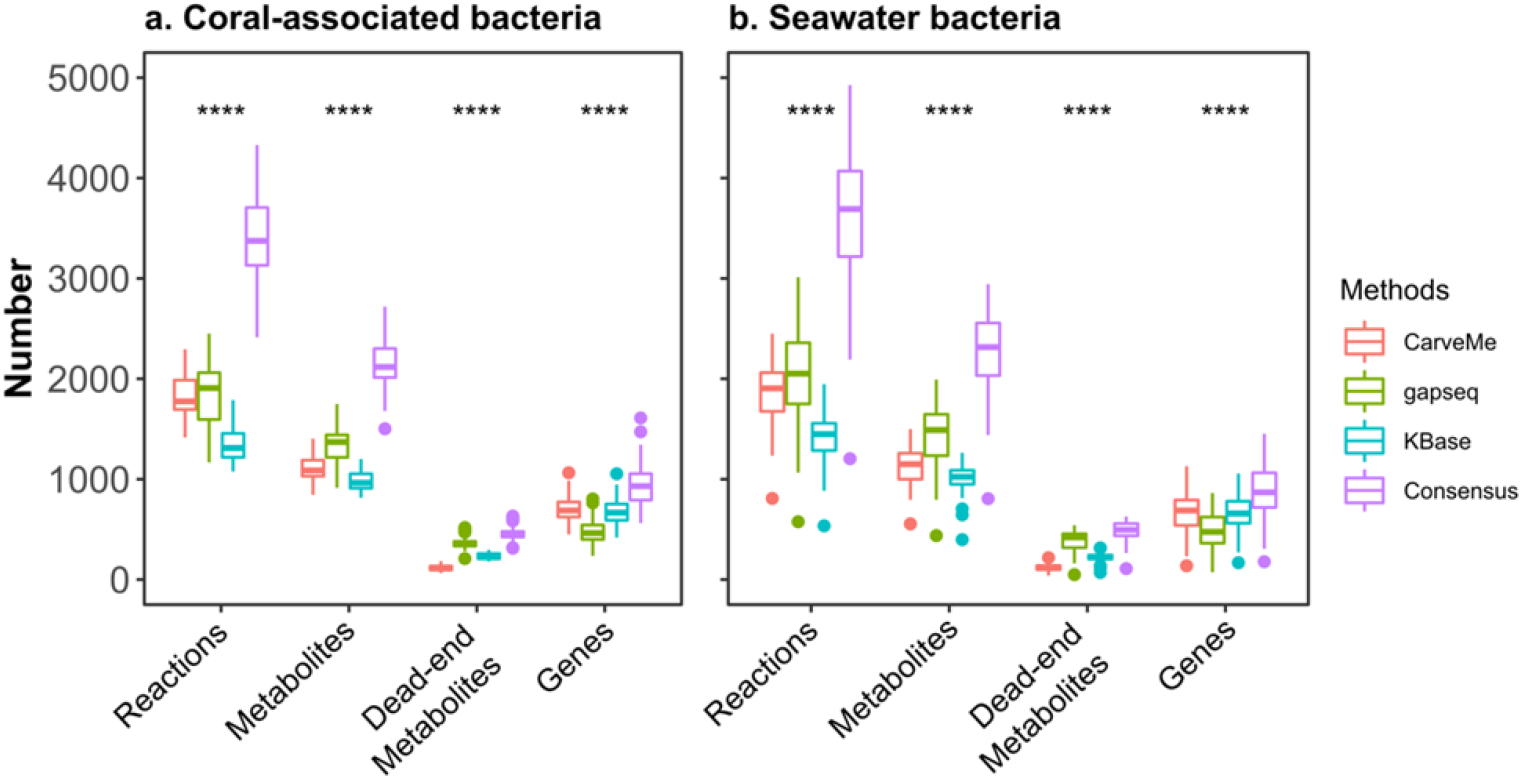
Structural comparison of metabolic models reconstructed using four different approaches. The metabolic models reconstructed by four approaches, including: CarveMe (16), gapseq (17), KBase (18), and the consensus method used in COMMIT (11), were evaluated based on the number of reactions, metabolites, dead-end metabolites, and genes. Statistical analysis was conducted using the Kruskal-Wallis test (**** p < 0.0001) to determine significant differences of these characteristics between methods. a. metabolic models of 50 coral-associated bacteria b. metabolic models of 55 seawater bacteria, based on MAGs from Robbins et al. (26). Each color represents a distinct reconstruction approach, as specified in the legend.

To assess the similarity of community reconstructions obtained through different approaches, we computed the Jaccard similarity for the sets of reactions, metabolites, dead-end metabolites, and genes in the models derived from the same MAGs (Fig. 2). Our findings revealed that despite being reconstructed from the same MAG, distinct reconstruction approaches yielded markedly different results. The results demonstrated a relatively low similarity between the respective sets resulting from the compared approaches. Specifically, in terms of the overall characteristics, gapseq and KBase models exhibited higher similarity in the composition of reactions and metabolites compared to CarveMe models. On average, the Jaccard similarity for reactions in coral-associated bacteria and seawater bacteria models was 0.23 and 0.24, respectively, while the Jaccard similarity for metabolites was 0.37 for models of both coral-associated and seawater bacterial communities. This observation suggests that the similarity between gapseq and KBase models may be attributed to their shared usage of the ModelSEED database for reconstruction, resulting in a relatively consistent set of reactions and metabolites within the models. However, in terms of gene composition, CarveMe and KBase models exhibited a higher degree of similarity compared to gapseq models. The average Jaccard similarity of the gene sets of coral-associated bacteria and seawater bacteria models was 0.42 and 0.45, respectively. Notably, we found a higher similarity between CarveMe and consensus models, with values of 0.75 and 0.77 for coral-associated bacteria and seawater bacteria models, respectively. This further indicated that the majority of genes included in the consensus models are due to their inclusion of CarveMe models.

**Fig 2.**
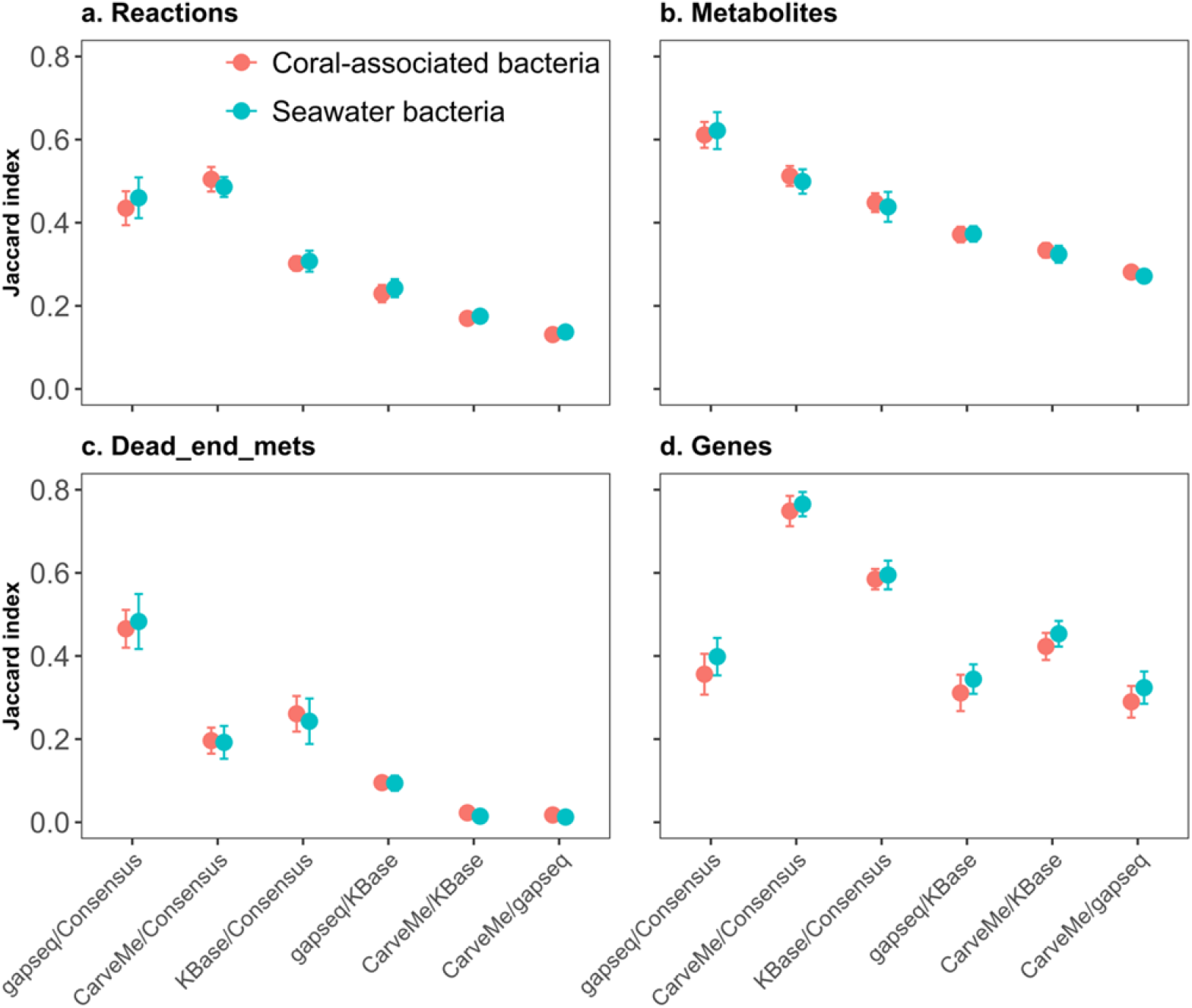
Analysis of similarity of community models derived from different reconstruction approaches. The Jaccard similarity was employed to assess the similarity between each reconstruction, considering: a. the sets of reactions, b. metabolites, c. dead-end metabolites, and d. genes. Pairwise comparisons were performed among the models reconstructed from the same MAG using different approaches. This comparison was performed on the same models whose characteristics were compared in Fig 1. The coral-associated bacteria models are represented in red, while the seawater bacteria models are depicted in light blue.

### The effects of iterative order on the reconstructed network

During the gap-filling process of the consensus models, we employed an iterative approach based on MAG abundance to specify the ascending/descending order of inclusion of a MAG in the gap-filling step of COMMIT. To investigate whether the order had an impact on the resulting gap-filling solutions, we conducted an analysis to assess the association between MAG abundance and the obtained solutions. Our findings indicated that the iterative order did not have a significant influence on the number of added reactions in the two communities reconstructed using the four different approaches (Fig. 3a-d, Fig. S1a-d, Fig. S2a-d, and Fig. S3a-d). The results demonstrated that the number of added reactions and abundance of MAGs exhibited only a negligible correlation (r = 0 to 0.3). In addition, although gapseq models exhibiting a higher number of reactions compared to CarveMe and KBase models, a considerable number of reactions without genetic support needed to be added to enable simulation of growth with gapseq models (Figs. 4 and S4). This observation raised concerns regarding the potential impact on the accuracy of model predictions. In contrast, the consensus approach demonstrated its ability to significantly reduce the number of required gap-filling solutions, thus minimizing the inclusion of such reactions without genetic support that are necessary for growth simulation.

**Fig 3.**
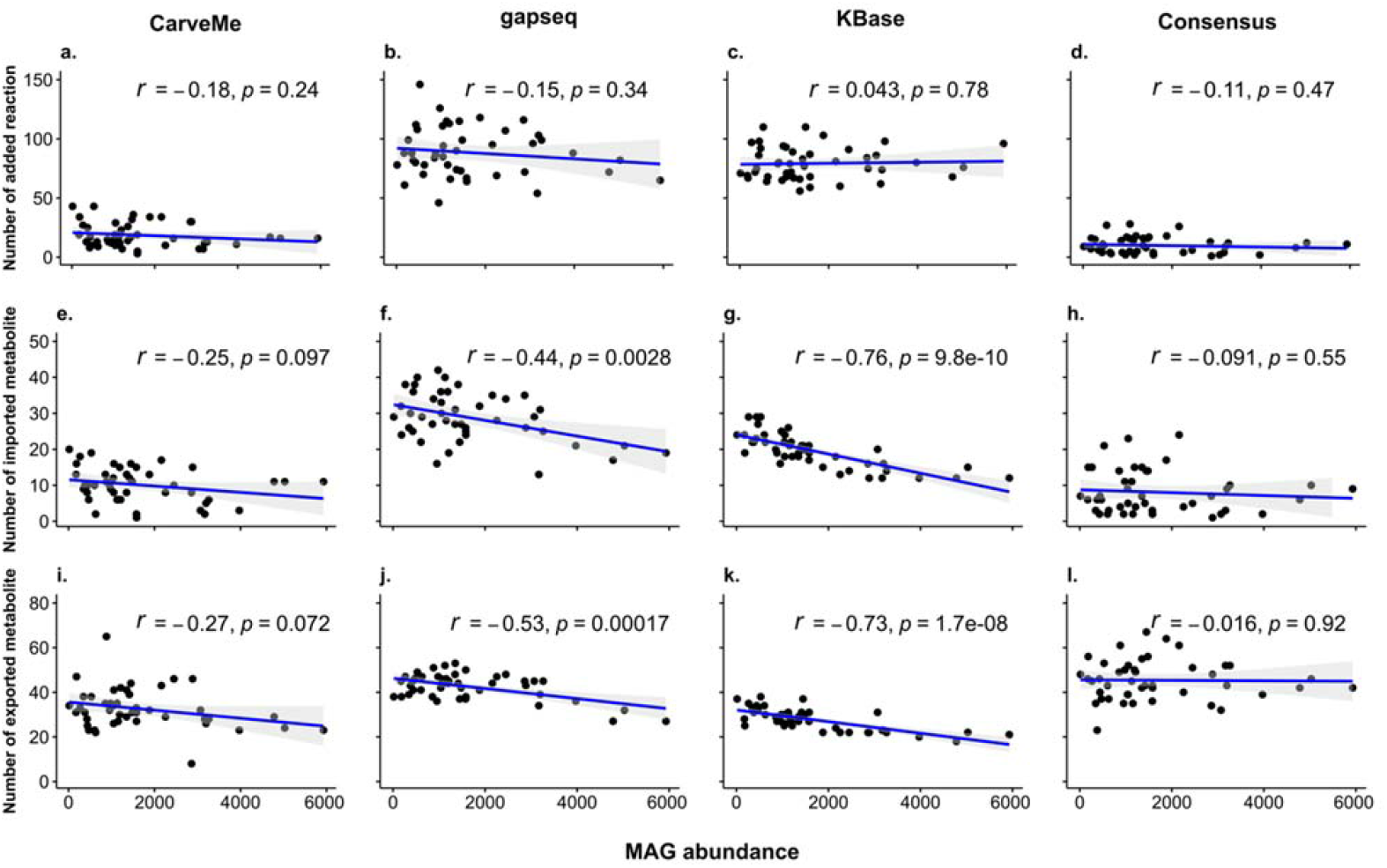
Association between MAG abundance and gap-filling results with a descending order in different reconstructions of coral-associated bacteria community model. Pearson correlation coefficient was employed to evaluate the association between MAG abundance and the number of added reactions (a - d), imported metabolites (e - h), and exported metabolites (i - l), for each of the four reconstruction approaches: CarveMe (16), gapseq (17), KBase (18), and the consensus method used in COMMIT (11). The correlation coefficient (*r*) and corresponding p-value (*p*) were determined.

**Fig 4.**
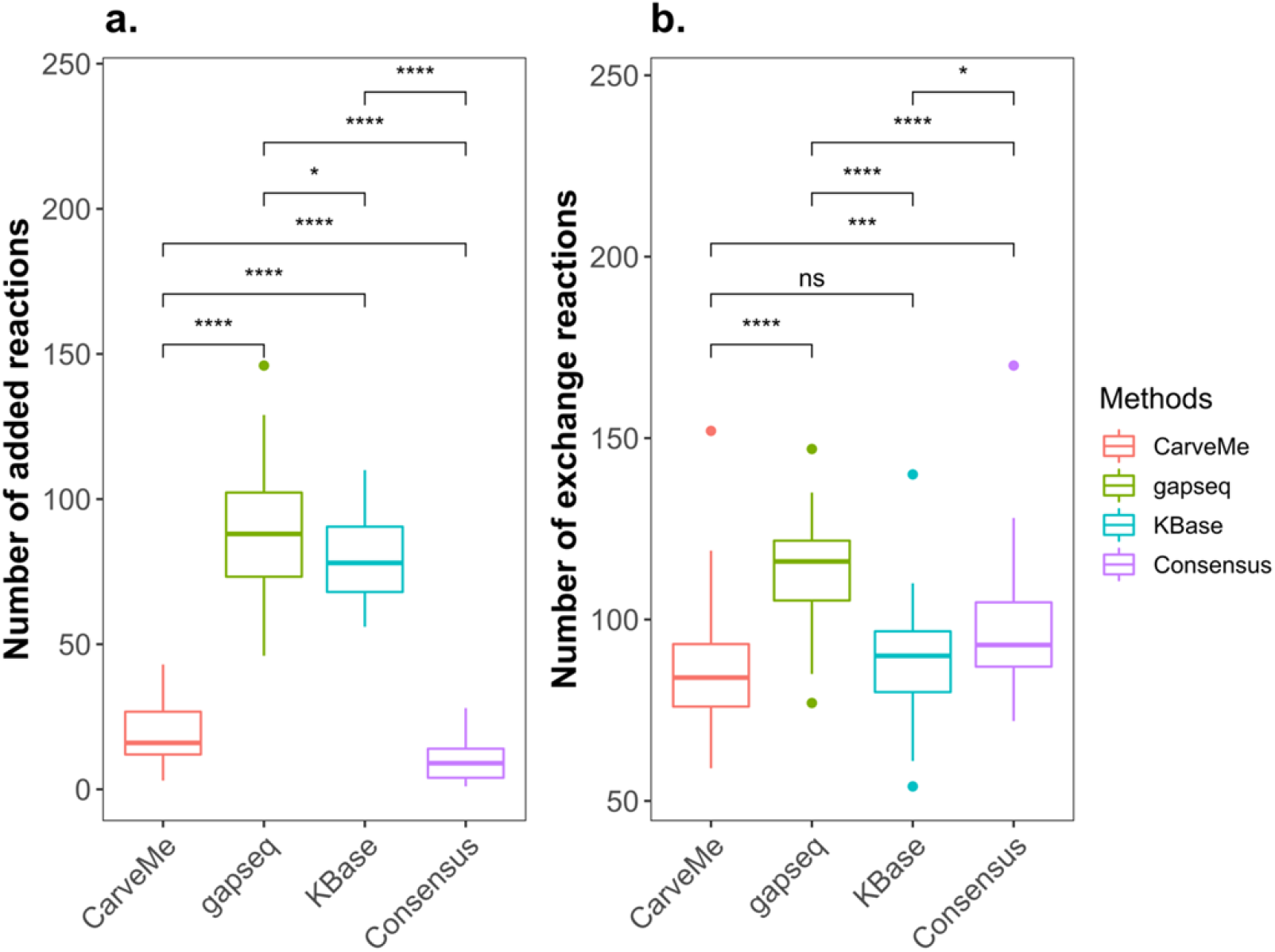
Comparison of functional models in different reconstructions of the coral-associated bacteria community model. The size of gap-filling solutions and the number of exchange reactions in functional models, that can simulate growth, were compared using the Wilcoxon Rank test (* p < 0.05; *** p < 0.001; **** p < 0.0001; ns p > 0.05). Panels a and b represent the size of gap-filling solutions and the number of exchange reactions, respectively.

With regards to the number of imported/exported metabolites (Figs. 3e-l, S1e-l, S2e-l, and S3e-l), the effect of MAG abundance in the order of iterative inclusion varied across different reconstruction approaches, with notable effects observed in the gapseq and KBase models. In CarveMe and consensus approaches, the MAG abundance order did not demonstrate a significant effect. In contract, for the KBase models we identified a high negative correlation (r = −0.7 to −0.9) between MAG abundance and the number of exported/imported metabolites (r = −0.76 and −0.73 for imported/exported metabolites, respectively). In the gapseq models, we found a low negative correlation (r = −0.3 to −0.5) between abundance and imported metabolites, while a moderate negative correlation (r = −0.5 to −0.7) existed between abundance and exported metabolites. However, when considering the increasing order of MAG abundance in KBase and gapseq models, the outcomes were reversed, demonstrating a positive correlation between MAG abundance and imported/exported metabolites. Regardless of the iterative order, it was noted that the starting model had a lower number of exchanged metabolites, while the ending model exhibited a higher number of exchanged metabolites in KBase and gapseq models. These findings suggest effects of reconstruction tools as well as abundance of MAGs on the exchange metabolites in the model.

### The quality assessment of functional models

Next, we performed an evaluation of the model quality using the MEMOTE suite of indices (Fig. 5). A higher score within this evaluation indicates better model quality according to the specified indices. The consistency index encompasses assessments of stoichiometric, mass, and charge balance of reactions, as well as metabolite connectivity and unbounded flux within the default medium. Notably, we stress the unbounded flux in the default medium index, as it elucidates the extent to which reactions can carry unlimited flux. This issue often arises due to problems with reaction directionality, missing cofactors, and/or inaccurately defined transport reactions (27). A higher score in this index correlates with a reduced number of reactions carrying unlimited flux. Another index we investigated is the reaction annotation index, which evaluates how many reactions in the model are annotated with associated EC numbers.

**Fig 5.**
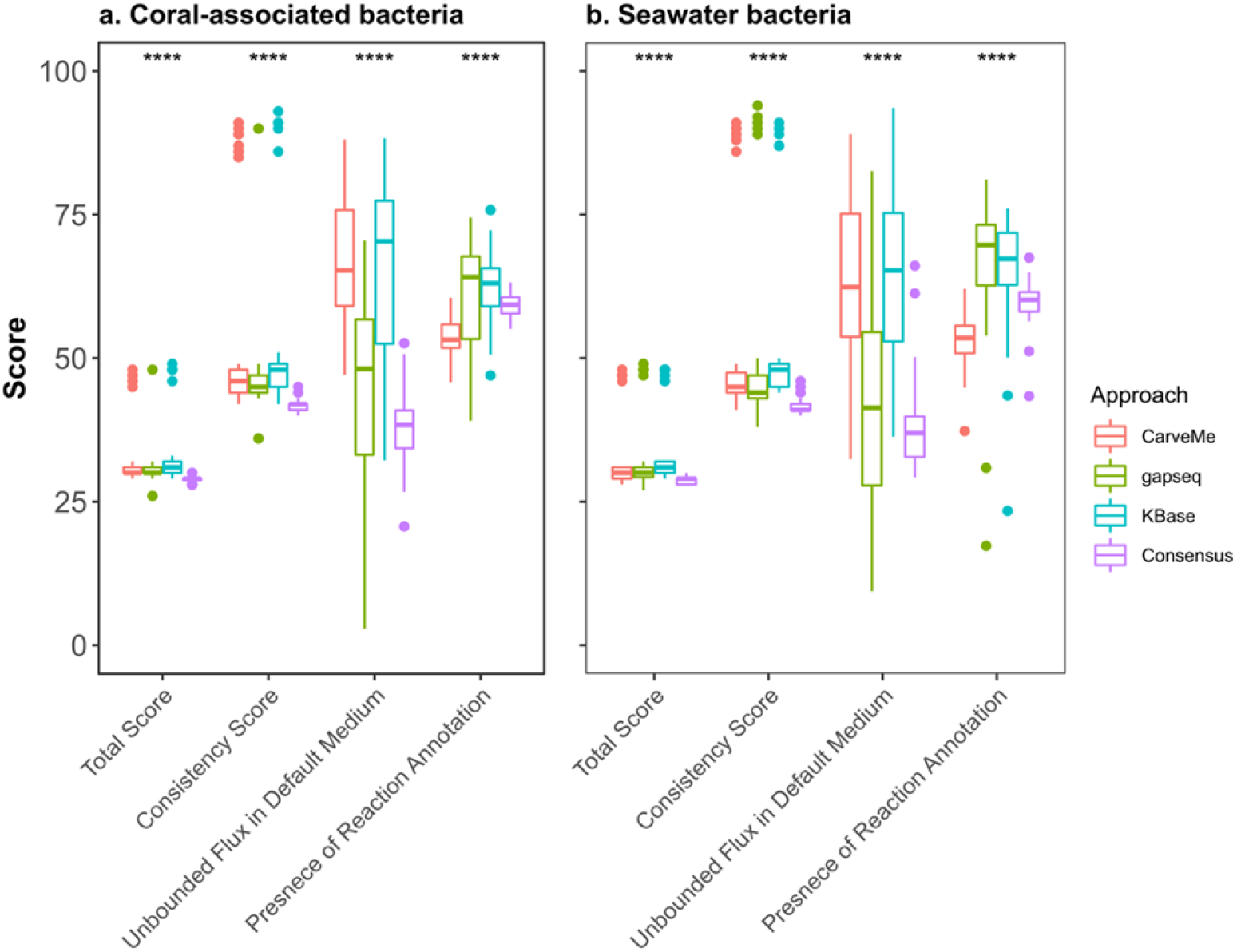
Quality assessment using MEMOTE. To assess the quality of models reconstructed from different approaches we used MEMOTE. Statistical analysis was conducted using the Kruskal-Wallis test (**** p < 0.0001) to determine significant differences of each score between methods. **a.** metabolic models of 50 coral-associated bacteria **b.** metabolic models of 55 seawater bacteria. Each color represents a distinct reconstruction approach, as specified in the legend.

We observed that the significant reduction in the total score was primarily attributed to the absence of reaction, metabolite, and gene annotations from databases other than MetaNetX. Regarding the individual scores, we found that KBase obtained the highest average score (62%) for the reaction annotation in the coral-associated bacteria models, while gapseq achieved the highest average score (67%) in the seawater bacteria models. Conversely, CarveMe exhibited the lowest score (54%) in both the coral-associated bacteria and seawater bacteria models. Interestingly, we found considerable variation in each score within the same reconstruction approach, indicating substantial differences in model quality. However, the consensus model demonstrated a noteworthy reduction in the variability of index values across models in comparison to the other approaches.

### Functional enrichment in different reconstructions

EC numbers provide the means to assess the enzyme functions included in a model in an automated fashion (28). For instance, enriched EC numbers can serve as an indicator of enriched function of metabolic reaction in a metabolic network.

To investigate the enriched functions in the reconstructed models, we performed a comparison of enriched EC numbers for the shared reactions and unblocked shared reactions in the models resulting from the compared approaches (Figs. 6a, S5a). The unblocked shared reactions were identified by doing flux variability analysis (FVA) among all shared reactions between the models. Although gapseq and KBase models exhibit relatively similar sets of reactions, our enrichment analysis revealed distinct enriched functions between these two approaches in terms of shared and unblocked shared reactions. For example, in the shared reactions within gapseq and KBase models, we observed an enrichment of functions related to acyltransferases and carbon-carbon lyases. However, after filtering blocked shared reactions, we found that glycosyltransferases and the enzymes involved in transferring nitrogenous groups and transferring phosphorus containing groups to be enriched. This discrepancy suggests that certain shared reactions in the gapseq and KBase models may not carry flux, thereby contributing to the observed differences.

**Fig 6.**
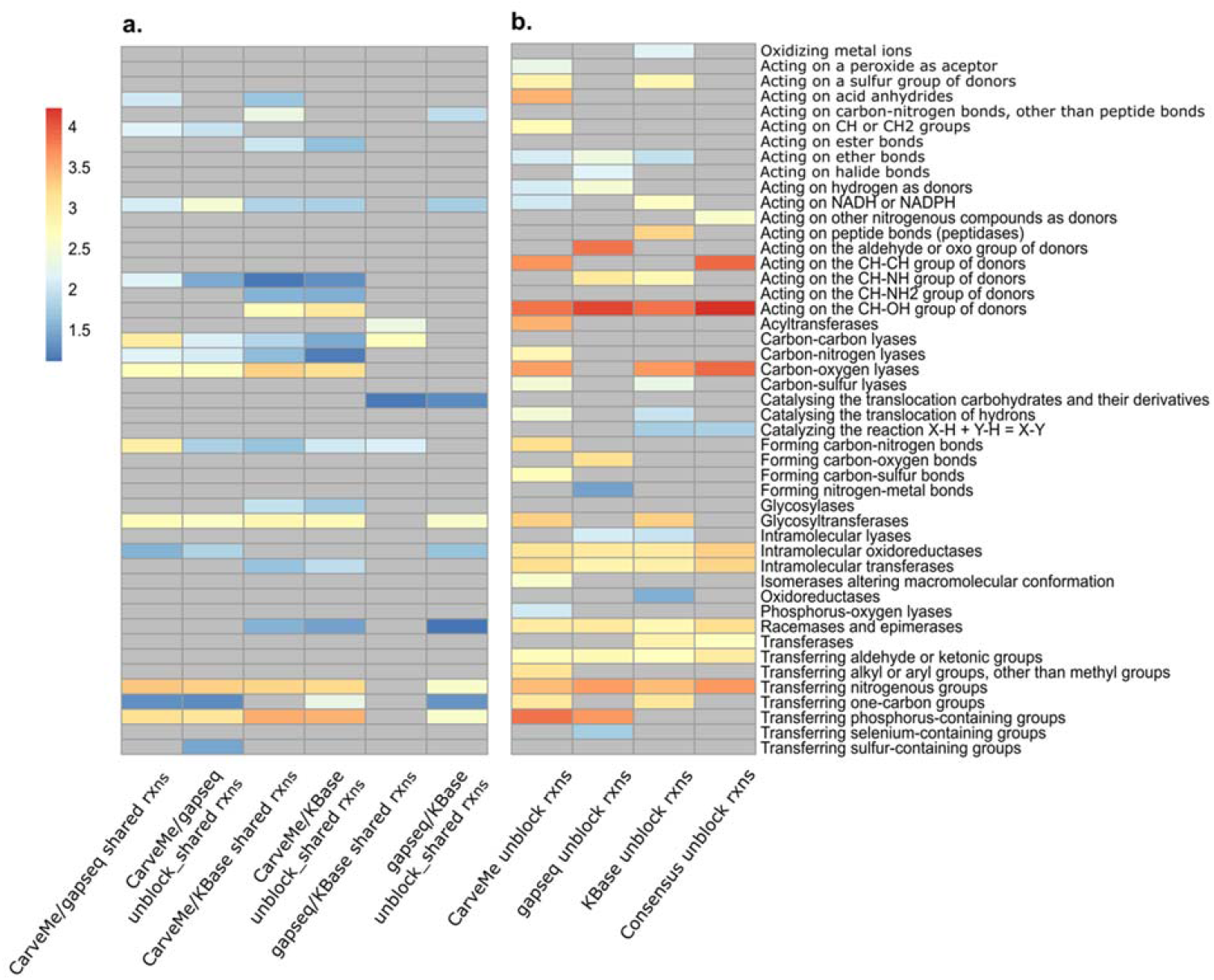
Enriched enzyme subclasses in the coral-associated bacteria community model from different reconstructions. The pairwise comparison of enriched enzyme subclasses in **a.** shared reactions between each reconstruction and **b.** in the community models reconstructed by different approaches, analyzed using the hypergeometric test. The abundance of enzyme subclasses is represented in a logarithmic scale and depicted using a color scale ranging from blue to red, with higher numbers indicating greater abundance. Grey color indicates the absence of enriched enzyme subclasses.

Conversely, we observed a higher degree of consistency in the enriched functions associated with shared and unblocked shared reactions in CarveMe/gapseq and CarveMe/KBase models. These consistent functions primarily encompassed activities related to carbon-oxygen lyases, glycosyltransferases, and the enzymes involved in transferring nitrogenous groups and transferring phosphorus containing groups. Overall, we found that CarveMe models displayed a greater diversity of enriched functions compared to gapseq and KBase models (Figs. 6b, S5b). The consensus models displayed more specific enriched functions. Predominantly enriched functions within both bacterial communities were associated with carbon-oxygen lyases and oxidoreductases, specifically those involved in acting on CH-OH and CH-CH group donors. This observation underscores the potential of consensus models to provide a more precise representation of the functional characteristics in bacterial community models. Overall, our results indicated that the seawater bacterial community displayed a higher diversity of enriched functions (13 enriched functions) compared to the coral-associated bacteria community (11 enriched functions).

### Exchanged metabolites in different reconstructions under community setting

We considered the presence of exchanged metabolites in the community models as a potential indicator of metabolite interactions. Sink reactions and exchange reactions were utilized within the community models to identify exported and imported metabolites, respectively. The intersection of these exported and imported metabolites constituted the set of exchanged metabolites, denoting metabolites that could be both secreted and taken up by members of the bacterial community. Our analysis revealed that consensus community models exhibited higher number of exported metabolites (Table 1). On average, each model secreted 44.8±9.1 and 42.8±6.9 metabolites within the coral-associated bacteria and seawater bacteria community, respectively. However, despite the large number of metabolites available for secretion into the medium within the community, only 64 metabolites were found to be exchanged within the community. The highest number of exchanged metabolites was observed in gapseq models for the coral-associated bacteria community (92 exchanged metabolites) and in CarveMe models for the seawater bacteria community (90 exchanged metabolites).

**Table 1.** The average number of imported, exported metabolites per model, and the number of exchanged metabolites in the community from different reconstruction approaches.

|  | Methods | Imported<br>metabolites | Exported<br>metabolites | Exchanged<br>metabolites |
| --- | --- | --- | --- | --- |
| **Coral-associated**<br>**bacteria** | CarveMe | 10.7±7.1 | 31.7±9.1 | 80 |
|  | gapseq | 28.1±8.1 | 40.7±7.8 | 92 |
|  | KBase | 19.1±7.2 | 26.4±6.4 | 49 |
|  | Consensus | 8.6±8.1 | 44.8±9.1 | 64 |
| **Seawater bacteria** | CarveMe | 10.4±7.6 | 29±6.9 | 90 |
|  | gapseq | 24.1±7.8 | 38.1±5.9 | 87 |
|  | KBase | 19.2±7.5 | 27.2±6.9 | 50 |
|  | Consensus | 7.4±8.7 | 42.8±6.9 | 64 |

Regarding the similarity of exchanged metabolites (Fig S6), the gapseq and KBase models exhibited relatively similar sets of exchanged metabolites compared to the CarveMe models in both the coral-associated and seawater bacterial communities (Jaccard index of 0.34 in both communities). This finding suggests that the use of the same database for model reconstruction may contribute to the similarity in exchanged metabolites among these approaches. Furthermore, our results indicate that the types of exchanged metabolites within the community models are highly dependent on the chosen reconstruction approaches and the underlying databases. Interestingly, community models reconstructed using the same approach, even if applied to different communities, displayed more similar sets of exchanged metabolites compared to community models reconstructed using different approaches. This finding warrants careful consideration of the conclusions drawn from applications of these models to assess the functional relevance of microbial interactions in communities.

## Discussion

In this study, we employed both top-down and bottom-up approaches for reconstruction of community models on the test case of coral-associated and seawater bacterial communities. The resulting models were subsequently compared with the consensus community models. To minimize the inherent uncertainty associated with each approach, we maintained uniformity by utilizing the same gene annotation tool (RAST) and adopting a universal biomass reaction during the model reconstruction process. However, despite these standardized procedures, we found substantial structural disparities among the resulting community models. We attribute these variations primarily to the gene-reaction mapping in the employed databases, which can significantly impact the model outcomes.

Gene sets are the basis of reconstructing GEMs. The absence of a gene in a model can result from the unavailability of its orthologous gene in the database or a lack of associated reactions within the database. It is generally assumed that models sharing similar gene sets would also exhibit similarity in their sets of reactions. However, CarveMe and KBase models demonstrated contradictory outcomes in terms of the similarity between gene and reaction sets. This finding could be attributed to differences in gene-reaction association information present in the BiGG and ModelSEED databases. This may also be a result of the variation in the number of reactions between CarveMe and KBase models. Additionally, the number of genes does not show positive correlation with the number of reactions or the proportion of reactions supported by genetic evidence. While gapseq models showcased a comparatively smaller number of genes, they encompassed a significant number of reactions, with merely 7.7% and 8.3% of total reactions on average lacking GPR associations in coral-associated and seawater bacterial models, respectively. This divergence might be attributed to the use of a customized database within the gapseq approach, which seemingly provided more comprehensive information regarding gene-reaction associations and resulting in numerous genes being associated with multiple reactions.

In this study, we also applied FVA to identify and filter out blocked reactions within the models, allowing us to investigate the enriched functions in active reactions in the community models. We observed that acyltransferases, which participate in the synthesis of long-chain fatty acids (29), was enriched in the shared reactions of gapseq and KBase models. However, this enrichment was not observed in the unblocked shared reactions. We hypothesize that the same reactions may carry different fluxes in the models reconstructed from different approaches, which can influence the enriched functions of models. Interestingly, the consensus approach demonstrated a greater capacity to distinguish the functional characteristics of different community models. This may be attributed to the comprehensive representation of biochemical reactions in consensus models. For instance, enzymes with oxidoreductase activity, acting on X-H and Y-H to form an X-Y bond, with oxygen as acceptor and those transferring aldehyde or ketonic groups were exclusively enriched in the coral-associated bacterial community. Conversely, enzymes associated with carbon-sulfur lyases, acting on ether bonds, acting on sulfur group donors, as well as acyltransferases were specific to the seawater bacterial community.

COMMIT employs a costless secretion approach to construct the community model. This iterative process allows us to simulate microbial interactions in terms of metabolite exchange and significantly reduces the number of added reactions required for the community networks (11). The order of iteration appears to have minimal influence on the number of added reactions across different reconstruction approaches. Nonetheless, in gapseq and KBase models, we observed a negative correlation between the number of imported/exported metabolites and the MAG abundance. Remarkably opposite outcomes were encountered when we applied the iterative process using MAG abundance in increasing order for gapseq and KBase models. These findings underscore the potential impact of the iterative process on the metabolite secretion/take-up capacity of models reconstructed from gapseq and KBase. In contrast, CarveMe and consensus models exhibited no correlation with the iterative order, which might help to mitigate uncertainties related to the metabolite transport capabilities of the models.

Costless metabolites have been instrumental in studying interspecies interactions in microbial ecosystems and have been suggested as mechanisms for maintaining genetic diversity within communities (30). Hence, in this study, we also examined the exchanged metabolites within the community models. We hypothesized that different reconstruction approaches would present distinct interaction outcomes. We observed a considerable variation in the number of exported/imported metabolites per model under different reconstruction approaches, leading to differences in the count of metabolites that could be exchanged within the community. Notably, our findings highlighted that the profile of exchanged metabolites was more influenced by the reconstruction approach used than by the type of bacterial community. This observation suggests a potential bias in predicting metabolite interactions using community GEMs. To mitigate such biases and improve the accuracy of phenotypic predictions from community models, the integration of additional information, such as omics data, is necessary.

Overall, the consensus approach effectively integrates a majority of the information derived from diverse reconstruction tools into a unified model. For example, the consensus model incorporates all genes present in the models reconstructed from the same MAG, along with a substantial number of reactions and metabolites. This integration leads to a notable reduction in the required gap-filling solutions. However, during this incorporation process, the consensus models also may assimilate all the dead-end metabolites and unbounded flux reactions inherent in the original models. This assimilation, in turn, may result in a reduction in the quality of the model. Despite this potential drawback, we found that the consensus approach results in good quality of models which is shared by the majority of models in the community–a feature which is not typical for the other approaches. By mitigating the variability of model quality, the consensus approach may potentially lead to better prediction of exchange metabolites in the bacterial community.

## Material and Methods

### Metagenome-assembled genome data

A total of 105 bacterial metagenome-assembled genomes (MAGs) were downloaded from NCBI under Bioproject accession PRJNA545004 (26) to reconstruct community metabolic models. Altogether 50 MAGs were from the coral tissue of *Porites lutea*, including the bacteria phyla: Acidobacteriota (6 MAGs), Actinobacteriota (2 MAGs), Bacteroidota (3 MAGs), Chloroflexota (14 MAGs), Dadabacteria (1 MAG), Gemmatimonadota (4 MAGs), UBP10 (2 MAGs), Latescibacterota (3 MAGs), Nitrospirota (1 MAG), Poribacteria (7 MAGs), and Proteobacteria (7 MAGs). These phyla broadly represent the taxonomic diversity observed in *P. lutea* (26). In addition, 55 MAGs were used to represent the bacteria phyla composition of surrounding seawater around Orpheus Island, Great Barrier Reef, Australia which harbours: Actinobacteriota (3 MAGs), Bacteroidota (10 MAGs), Cyanobacteriota (2 MAGs), Marinisomatota (1 MAG), Patescibacteria (1 MAG), Planctomycetota (4 MAGs), Proteobacteria (30 MAGs), SAR324 (2 MAGs), and Verrucomicrobiota (2 MAGs). Abundance of MAGs was determined using BBMap (31), which calculated the average read coverage of per contigs per MAG, generating a coverage profile. The abundance of MAGs was then presented as the sum of contig coverages.

### Generation of draft and consensus metabolic models

Metabolic reconstruction approaches rely on diverse databases, and the choice between bottom-up and top-down methodologies can lead to variations in the structure of metabolic reconstructions. To provide a comprehensive overview of this discrepancy, we compared three reconstruction approaches: CarveMe (16), gapseq (17), and KBase (18). Among these approaches, CarveMe belongs to top-down reocnstruction approach while gapseq and KBase are bottom-up approaches. For the reconstruction, the MAGs were annotated using Annotate Metagenome Assembly and Re-annotate Metagenome with RASTtk – v1.073 app (32–34) published on KBase platform.

In the CarveMe reconstruction approach, the manually curated universal bacteria model was used as a template. The annotated sequence was aligned with the amino sequences in the BiGG database (35). Subsequently, the reaction scores were derived by associating them with the sequence similarity scores through the gene-protein reaction (GPR) rules. Reactions lacking genetic evidence were assigned negative scores within this framework. During the model carving process, reactions with low scores were eliminated from the universal model to generate the draft models.

In the gapseq tool, draft models were reconstructed using the default settings. The annotated sequences were utilized to predict pathways and subsystem using a customized database, obtained from MetaCyc (36), KEGG (37), and ModelSEED (38). Additionally, transporters were predicted based on the Transporter Classification Database (TCDB) (39), which catalogs a wide range of transport proteins and their functional classifications.

In the KBase approach, the metabolic reconstruction process was carried-out using the ModelSEED pipeline (38). The functional annotation of MAGs obtained from RAST was directly mapped to the corresponding biochemical reactions present in the ModelSEED biochemistry database. The biomass reactions were based on a template biomass reaction and assigned non-universal biomass components, such as cofactors and cell wall components, using the SEED subsystems and RAST functional annotations. Subsequently, the draft models resulting from this process were downloaded for further analysis and refinement.

To build the consensus models, we followed the pipeline provided in COMMIT (11). Before merging the models obtained from different reconstructions, we unified the reaction and metabolite IDs in the draft models by mapping them to MNXref IDs using the provided MNXref reference files (40). The biomass reaction, if present, and exchange reactions were subsequently removed. In the merging process, we used the CarveMe models as the initial component of the consensus models in an iterative fashion (following by gapseq and KBase models). Subsequent reconstructions were compared to this consensus model in a stepwise manner. First, the fields of the models were harmonized to ensure consistency. Next, the gene identifiers were compared, and if necessary, any genes not present in the consensus model were added. Subsequently, the reactions were compared based on various criteria, including reaction IDs, GPR rules, metabolite composition, reversibility, and mass balance. Any duplicate reactions and metabolites were removed from the consensus model to avoid redundancies.

### Gap-fill community models obtained by COMMIT

Before the gap-filling, the exchange and biomass reactions were removed from the draft CarveMe, gapseq, and KBase models. Subsequently, a universal biomass reaction, which was adapted from *Escherichia coli* biomass composition (41) according to the universal biomass components in prokaryotes (42), was added into the draft CarveMe, gapseq, KBase, and consensus models. To perform the iterative gap-filling, the community models derived from different reconstructions were processed in descending and acesding order given by the species abundance. Initially, a common microbial growth medium (LB media) was provided as the initial media for the gap-filling process. Adjusted M9 media (with glucose and magnesium ion) was then employed for subsequent iterations.

### Comparison of community models from different reconstructions

The quality of the models was assessed using MEMOTE (27) to evaluate their overall performance. In addition, several model features, including the number of reactions, metabolites, dead-end metabolites, and genes, were analyzed to compare the structural properties of the models. The similarity between models was determined using the Jaccard similarity coefficient.

To identify enriched functions in the community models, we extracted the enzyme commission numbers (EC numbers) from each reaction and used them to the second digit (i.e. enzyme subclass level) in the enrichment analysis. To this end, we conducted a hypergeometric test to identify significantly enriched EC numbers in the shared (unblocked) reactions between two models reconstructed from the same MAG using different approaches.

To identify potential exchange metabolites in the community, we examined the sink and exchange reactions in models, which allowed us to identify the metabolites involved in exchange processes. We also considered the exchanged metabolites as an indicator of metabolic interaction potential, enabling the evaluation of the metabolic interactions within the community models.

## Supplemental Material

**Fig S1.**
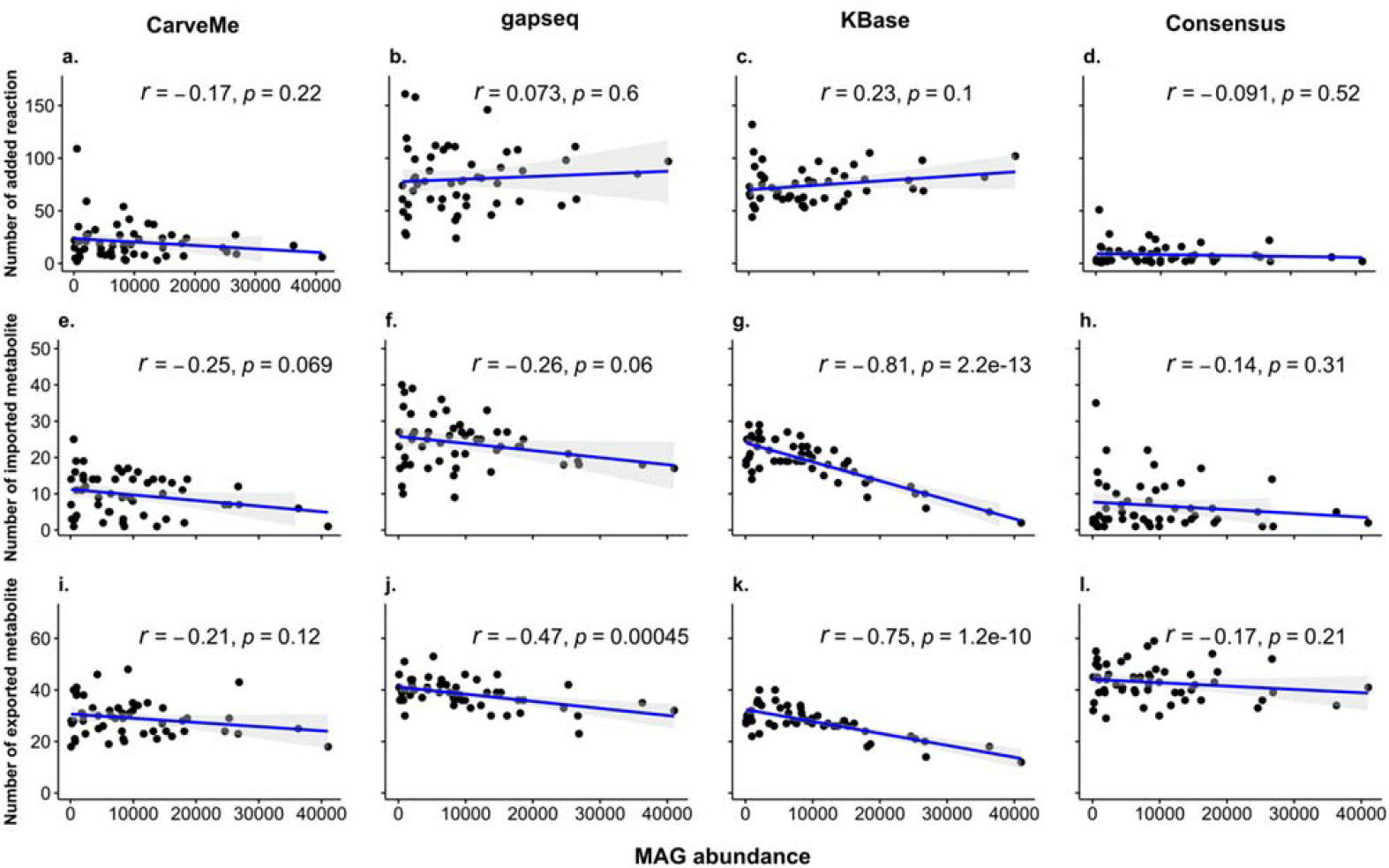
Association between MAG abundance and gap-filling results a descending order in different reconstructions of seawater bacteria community model. Pearson correlation coefficient was employed to evaluate the association between MAG abundance and the number of added reactions (a - d), imported metabolites (e - h), and exported metabolites (i - l), for each of the four reconstruction approaches: CarveMe (16), gapseq (17), KBase (18), and the consensus method used in COMMIT (11). The correlation coefficient (*r*) and corresponding p-value (*p*) were determined.

**Fig S2.**
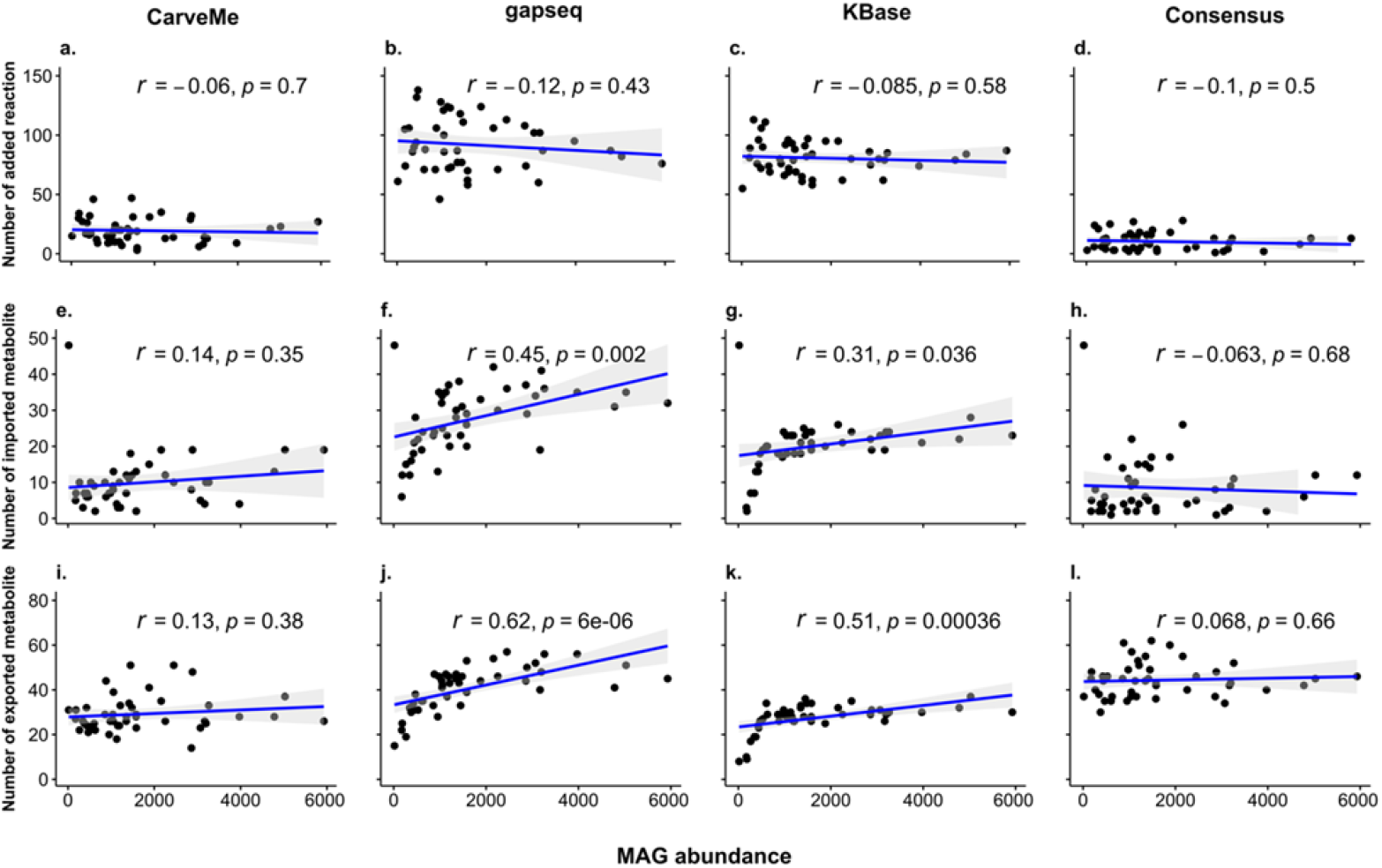
Association between MAG abundance and gap-filling results with an ascending order of MAG abundance in different reconstructions of coral-associated bacteria community model. Pearson correlation coefficient was employed to evaluate the association between MAG abundance and the number of added reactions (a - d), imported metabolites (e- h), and exported metabolites (i - l), for each of the four reconstruction approaches: CarveMe (16), gapseq (17), KBase (18), and the consensus method used in COMMIT (11). The correlation coefficient (*r*) and corresponding p-value (*p*) were determined.

**Fig S3.**
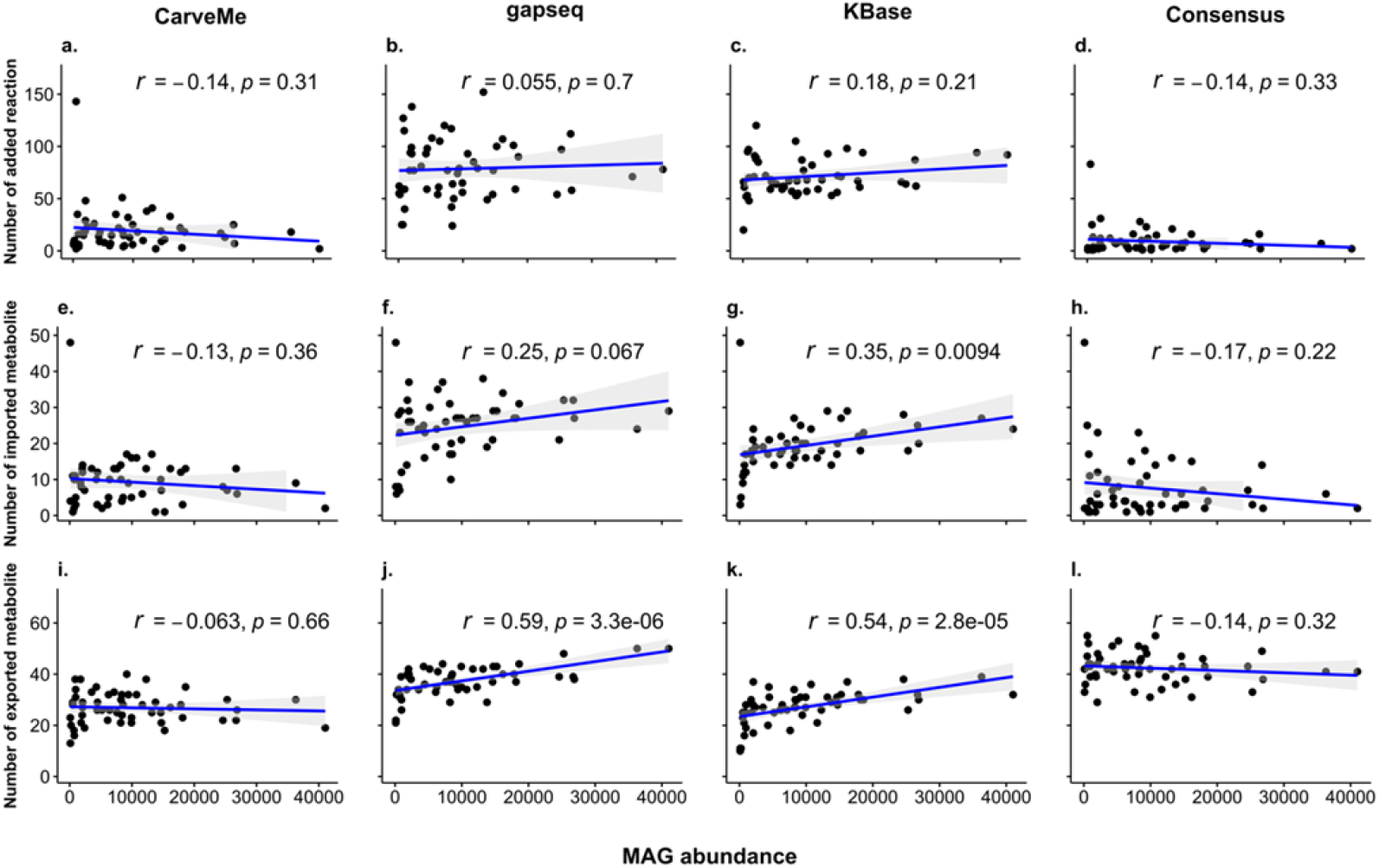
Association between MAG abundance and gap-filling results with an ascending order of MAG abundance in different reconstructions of seawater bacteria community model. Pearson correlation coefficient was employed to evaluate the association between MAG abundance and the number of added reactions (a - d), imported metabolites (e - h), and exported metabolites (i - l), for each of the four reconstruction approaches: CarveMe (16), gapseq (17), KBase (18), and the consensus method used in COMMIT (11). The correlation coefficient (*r*) and corresponding p-value (*p*) were determined.

**Fig S4.**
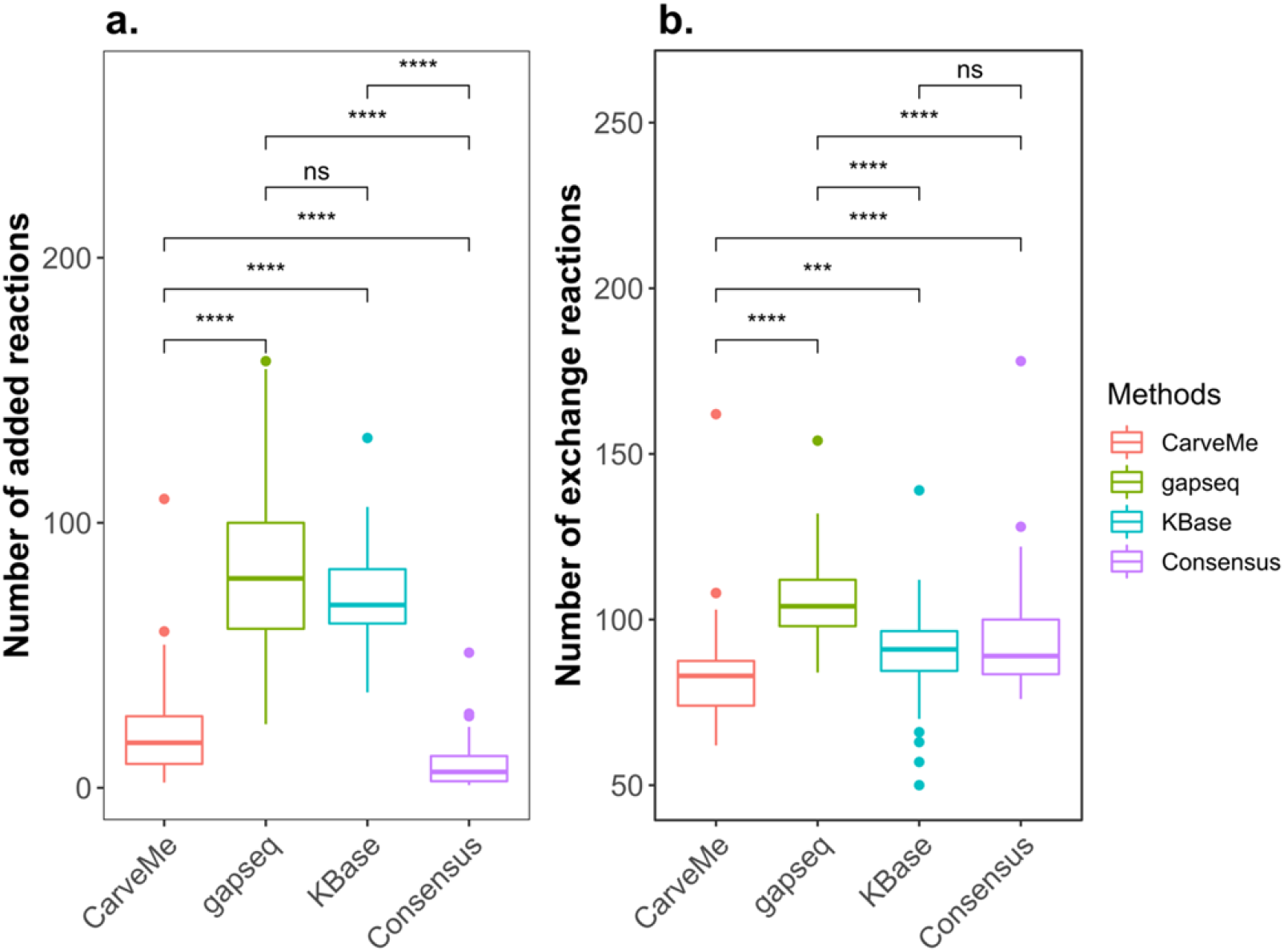
Comparison of functional models in different reconstructions of the seawater bacteria community model. The size of gap-filling solutions and the number of exchange reactions in functional models, that can simulate growth, were compared using the Wilcoxon Rank test (* p < 0.05; *** p < 0.001; **** p < 0.0001; ns p > 0.05). Panels a and b represent the size of gap-filling solutions and the number of exchange reactions, respectively.

**Fig S5.**
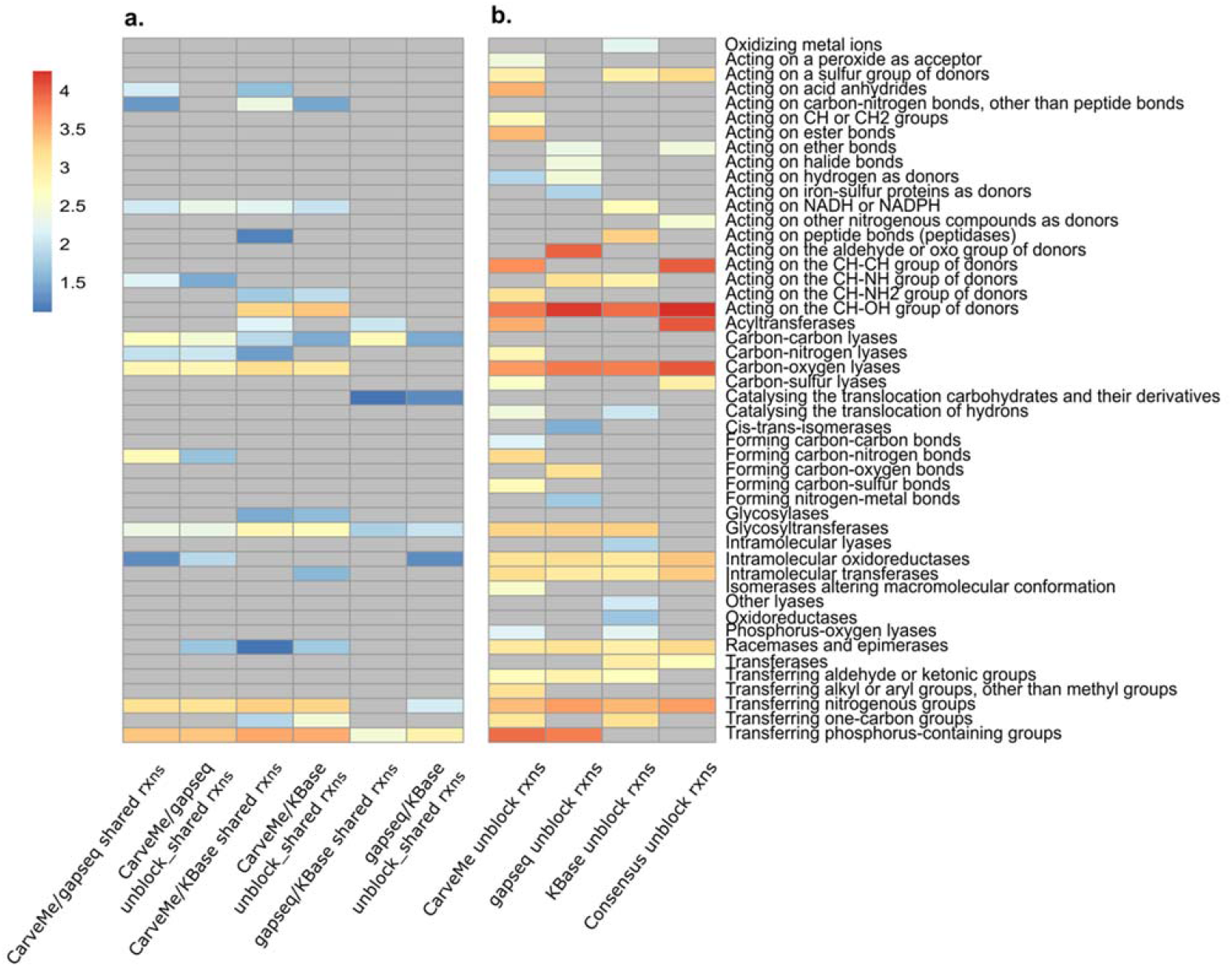
Enriched enzyme subclasses in the seawater bacteria community model from different reconstructions. The pairwise comparison of enriched enzyme subclasses in a. shared reactions between each reconstruction and b. in the community models reconstructed by different approaches, analyzed using the hypergeometric test. The abundance of enzyme subclasses is represented in a logarithmic scale and depicted using a color scale ranging from blue to red, with higher numbers indicating greater abundance. Grey color indicates the absence of enriched enzyme subclasses.

**Fig S6.**
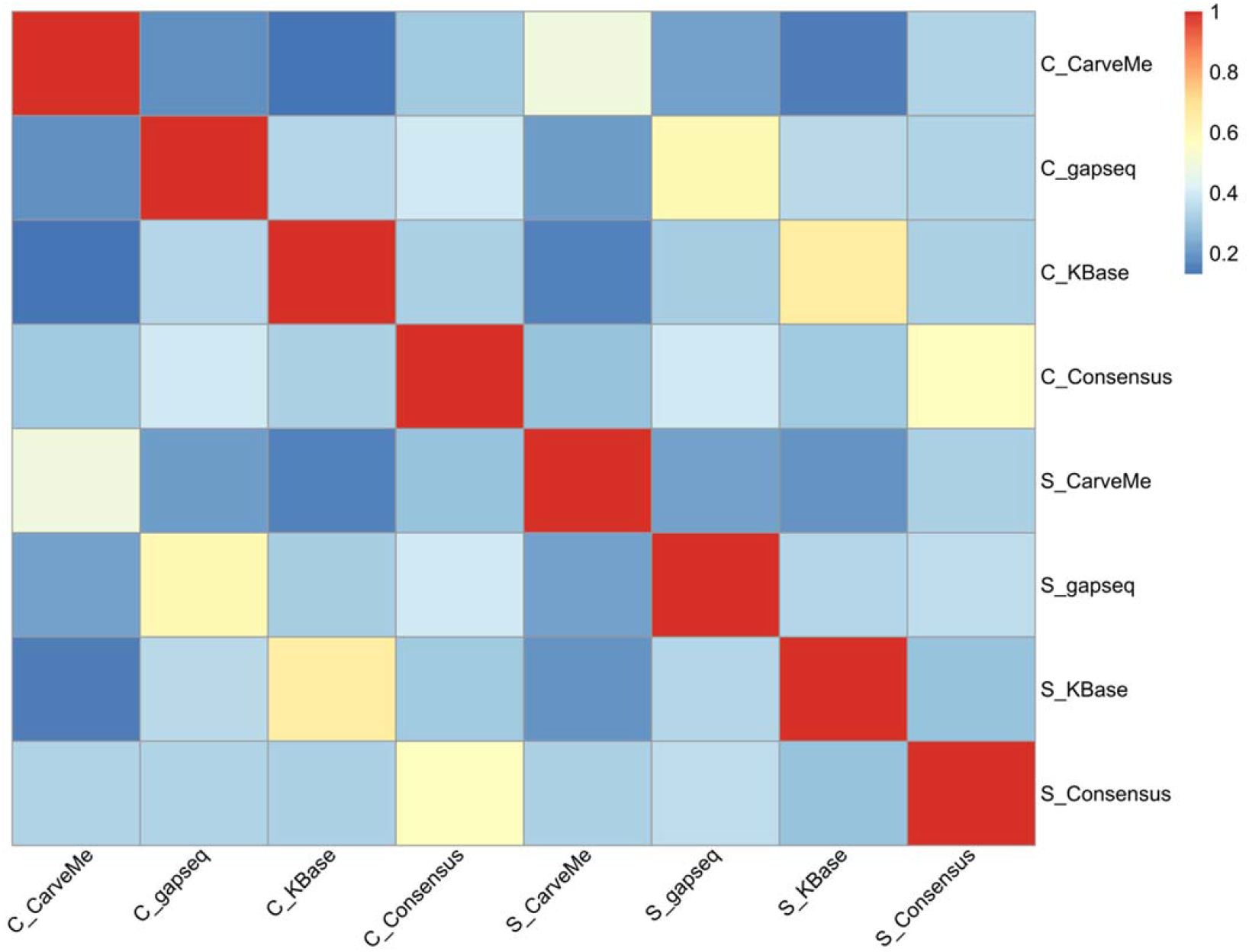
Jaccard index matrix of exchanged metabolites between different reconstruction approaches across the coral-associated and seawater bacteria communities. The pairwise comparison of exchanged metabolites within the community, derived from different approaches, was assessed using the Jaccard similarity. The labels ’C_’ and ’S_’ correspond to the coral-associated bacteria and seawater bacteria communities, respectively. The Jaccard index, ranging from 0 to 1, is visualized using a color scale that transitions from blue to red.

**Table S1.**
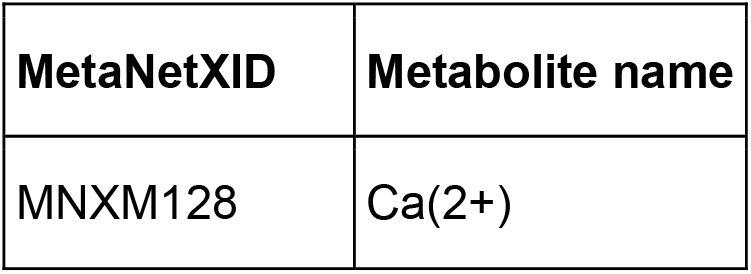
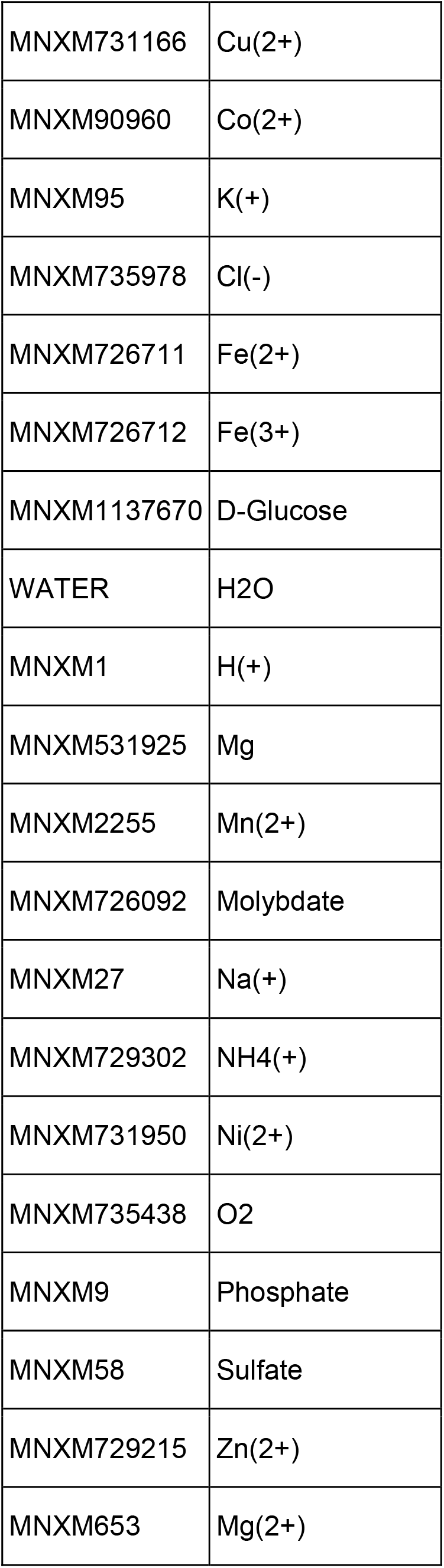
The M9 medium used for gap-filling.

## Acknowledgments

The authors would like to thank the Melbourne-Potsdam PhD Program of the Max Planck Institute of Molecular Plant Physiology, Potsdam, Germany and The University of Melbourne, Parkville, Melbourne for supporting this project.

## Reference

1. Konopka A, Lindemann S, Fredrickson J. 2015. Dynamics in microbial communities: unraveling mechanisms to identify principles. The ISME Journal 9:1488–1495.

2. Lawson CE, Wu S, Bhattacharjee AS, Hamilton JJ, McMahon KD, Goel R, Noguera DR. 2017. Metabolic network analysis reveals microbial community interactions in anammox granules. Nature Communications 8:15416.

3. Wang C-W, Wong J-WM, Yeh S-S, Hsieh YE, Tseng C-H, Yang S-H, Tang S-L. 2022. Soil Bacterial Community May Offer Solutions for Ginger Cultivation. Microbiology Spectrum 10:e01803–22.

4. De Roy K, Marzorati M, Van Den Abbeele P, Van De Wiele T, Boon N. 2014. Synthetic microbial ecosystems: an exciting tool to understand and apply microbial communities. Environmental Microbiology 16:1472–1481.

5. Althani AA, Marei HE, Hamdi WS, Nasrallah GK, El Zowalaty ME, Al Khodor S, Al-Asmakh M, Abdel-Aziz H, Cenciarelli C. 2016. Human Microbiome and its Association With Health and Diseases. Journal of Cellular Physiology 231:1688–1694.

6. de Voogd NJ, Cleary DFR, Polónia ARM, Gomes NCM. 2015. Bacterial community composition and predicted functional ecology of sponges, sediment and seawater from the thousand islands reef complex, West Java, Indonesia. FEMS Microbiology Ecology 91.

7. Pham VHT, Kim J. 2012. Cultivation of unculturable soil bacteria. Trends in Biotechnology 30:475–484.

8. Magnúsdóttir S, Heinken A, Kutt L, Ravcheev DA, Bauer E, Noronha A, Greenhalgh K, Jäger C, Baginska J, Wilmes P, Fleming RMT, Thiele I. 2017. Generation of genome-scale metabolic reconstructions for 773 members of the human gut microbiota. Nature Biotechnology 35:81–89.

9. Kundu P, Ghosh A. 2023. Genome-scale community modeling for deciphering the inter-microbial metabolic interactions in fungus-farming termite gut microbiome. Computers in Biology and Medicine 154:106600.

10. Du H, Pan J, Zou D, Huang Y, Liu Y, Li M. 2022. Microbial active functional modules derived from network analysis and metabolic interactions decipher the complex microbiome assembly in mangrove sediments. Microbiome 10:224.

11. Wendering P, Nikoloski Z. 2022. COMMIT: Consideration of metabolite leakage and community composition improves microbial community reconstructions. PLOS Computational Biology 18:e1009906.

12. Mataigne V, Vannier N, Vandenkoornhuyse P, Hacquard S. 2022. Multi-genome metabolic modeling predicts functional inter-dependencies in the Arabidopsis root microbiome. Microbiome 10:217.

13. Henry CS, Bernstein HC, Weisenhorn P, Taylor RC, Lee JY, Zucker J, Song HS. 2016. Microbial Community Metabolic Modeling: A Community Data-Driven Network Reconstruction. Journal of Cellular Physiology 231:2339–2345.

14. Gelbach PE, Finley SD. 2023. Flux Sampling in Genome-scale Metabolic Modeling of Microbial Communities. bioRxiv doi:10.1101/2023.04.18.537368.

15. Ang KS, Lakshmanan M, Lee NR, Lee DY. 2018. Metabolic Modeling of Microbial Community Interactions for Health, Environmental and Biotechnological Applications. Curr Genomics 19:712–722.

16. Machado D, Andrejev S, Tramontano M, Patil KR. 2018. Fast automated reconstruction of genome-scale metabolic models for microbial species and communities. Nucleic Acids Research 46:7542–7553.

17. Zimmermann J, Kaleta C, Waschina S. 2021. gapseq: informed prediction of bacterial metabolic pathways and reconstruction of accurate metabolic models. Genome Biology 22.

18. Arkin AP, Cottingham RW, Henry CS, Harris NL, Stevens RL, Maslov S, Dehal P, Ware D, Perez F, Canon S, Sneddon MW, Henderson ML, Riehl WJ, Murphy-Olson D, Chan SY, Kamimura RT, Kumari S, Drake MM, Brettin TS, Glass EM, Chivian D, Gunter D, Weston DJ, Allen BH, Baumohl J, Best AA, Bowen B, Brenner SE, Bun CC, Chandonia J-M, Chia J-M, Colasanti R, Conrad N, Davis JJ, Davison BH, DeJongh M, Devoid S, Dietrich E, Dubchak I, Edirisinghe JN, Fang G, Faria JP, Frybarger PM, Gerlach W, Gerstein M, Greiner A, Gurtowski J, Haun HL, He F, Jain R, et al. 2018. KBase: The United States Department of Energy Systems Biology Knowledgebase. Nature Biotechnology 36:566–569.

19. Mendoza SN, Olivier BG, Molenaar D, Teusink B. 2019. A systematic assessment of current genome-scale metabolic reconstruction tools. Genome Biology 20:158.

20. Bernstein DB, Sulheim S, Almaas E, Segrè D. 2021. Addressing uncertainty in genome-scale metabolic model reconstruction and analysis. Genome Biology 22:64.

21. Schnoes AM, Brown SD, Dodevski I, Babbitt PC. 2009. Annotation Error in Public Databases: Misannotation of Molecular Function in Enzyme Superfamilies. PLoS Computational Biology 5:e1000605.

22. Lobb B, Tremblay BJ-M, Moreno-Hagelsieb G, Doxey AC. 2020. An assessment of genome annotation coverage across the bacterial tree of life. Microbial Genomics 6.

23. Pham N, Van Heck R, Van Dam J, Schaap P, Saccenti E, Suarez-Diez M. 2019. Consistency, Inconsistency, and Ambiguity of Metabolite Names in Biochemical Databases Used for Genome-Scale Metabolic Modelling. Metabolites 9:28.

24. Chindelevitch L, Stanley S, Hung D, Regev A, Berger B. 2012. MetaMerge: scaling up genome-scale metabolic reconstructions, with application to Mycobacterium tuberculosis. Genome Biology 13:R6.

25. Aung HW, Henry SA, Walker LP. 2013. Revising the Representation of Fatty Acid, Glycerolipid, and Glycerophospholipid Metabolism in the Consensus Model of Yeast Metabolism. Industrial Biotechnology 9:215–228.

26. Robbins SJ, Singleton CM, Chan CX, Messer LF, Geers AU, Ying H, Baker A, Bell SC, Morrow KM, Ragan MA, Miller DJ, Forêt S, Ball E, Beeden R, Berumen M, Aranda M, Ravasi T, Bongaerts P, Hoegh-Guldberg O, Cooke I, Leggat B, Sprungala S, Fitzgerald A, Shang C, Lundgren P, Fyffe T, Rubino F, Van Oppen M, Weynberg K, Robbins SJ, Singleton CM, Xin Chan C, Messer LF, Geers AU, Ying H, Baker A, Bell SC, Morrow KM, Ragan MA, Miller DJ, Foret S, Voolstra CR, Tyson GW, Bourne DG, Voolstra CR, Tyson GW, Bourne DG. 2019. A genomic view of the reef-building coral Porites lutea and its microbial symbionts. Nature Microbiology 4:2090–2100.

27. Lieven C, Beber ME, Olivier BG, Bergmann FT, Ataman M, Babaei P, Bartell JA, Blank LM, Chauhan S, Correia K, Diener C, Dräger A, Ebert BE, Edirisinghe JN, Faria JP, Feist AM, Fengos G, Fleming RMT, García-Jiménez B, Hatzimanikatis V, Van Helvoirt W, Henry CS, Hermjakob H, Herrgård MJ, Kaafarani A, Kim HU, King Z, Klamt S, Klipp E, Koehorst JJ, König M, Lakshmanan M, Lee D-Y, Lee SY, Lee S, Lewis NE, Liu F, Ma H, Machado D, Mahadevan R, Maia P, Mardinoglu A, Medlock GL, Monk JM, Nielsen J, Nielsen LK, Nogales J, Nookaew I, Palsson BO, Papin JA, et al. 2020. MEMOTE for standardized genome-scale metabolic model testing. Nature Biotechnology 38:272–276.

28. Tipton K, Boyce S. 2000. History of the enzyme nomenclature system. Bioinformatics 16:34–40.

29. Röttig A, Steinbüchel A. 2013. Acyltransferases in bacteria. Microbiol Mol Biol Rev 77:277–321.

30. Morris JJ. 2015. Black Queen evolution: the role of leakiness in structuring microbial communities. Trends in Genetics 31:475–482.

31. Bushnell B. 2014. BBMap: a fast, accurate, splice-aware aligner. Lawrence Berkeley National Lab.(LBNL), Berkeley, CA (United States),

32. Aziz RK, Bartels D, Best AA, DeJongh M, Disz T, Edwards RA, Formsma K, Gerdes S, Glass EM, Kubal M, Meyer F, Olsen GJ, Olson R, Osterman AL, Overbeek RA, McNeil LK, Paarmann D, Paczian T, Parrello B, Pusch GD, Reich C, Stevens R, Vassieva O, Vonstein V, Wilke A, Zagnitko O. 2008. The RAST Server: Rapid Annotations using Subsystems Technology. BMC Genomics 9:75.

33. Overbeek R, Olson R, Pusch GD, Olsen GJ, Davis JJ, Disz T, Edwards RA, Gerdes S, Parrello B, Shukla M, Vonstein V, Wattam AR, Xia F, Stevens R. 2013. The SEED and the Rapid Annotation of microbial genomes using Subsystems Technology (RAST). Nucleic Acids Research 42:D206–D214.

34. Brettin T, Davis JJ, Disz T, Edwards RA, Gerdes S, Olsen GJ, Olson R, Overbeek R, Parrello B, Pusch GD, Shukla M, Thomason JA, Stevens R, Vonstein V, Wattam AR, Xia F. 2015. RASTtk: A modular and extensible implementation of the RAST algorithm for building custom annotation pipelines and annotating batches of genomes. Scientific Reports 5:8365.

35. King ZA, Lu J, Dräger A, Miller P, Federowicz S, Lerman JA, Ebrahim A, Palsson BO, Lewis NE. 2015. BiGG Models: A platform for integrating, standardizing and sharing genome-scale models. Nucleic Acids Research 44:D515–D522.

36. Caspi R, Billington R, Keseler IM, Kothari A, Krummenacker M, Midford PE, Ong WK, Paley S, Subhraveti P, Karp PD. 2019. The MetaCyc database of metabolic pathways and enzymes - a 2019 update. Nucleic Acids Research 48:D445–D453.

37. Kanehisa M, Sato Y, Furumichi M, Morishima K, Tanabe M. 2018. New approach for understanding genome variations in KEGG. Nucleic Acids Research 47:D590–D595.

38. Henry CS, Dejongh M, Best AA, Frybarger PM, Linsay B, Stevens RL. 2010. High-throughput generation, optimization and analysis of genome-scale metabolic models. Nature Biotechnology 28:977–982.

39. Saier MH, Jr, Reddy VS, Tamang DG, Västermark Å. 2013. The Transporter Classification Database. Nucleic Acids Research 42:D251–D258.

40. Moretti S, Tran Van Du T, Mehl F, Ibberson M, Pagni M. 2020. MetaNetX/MNXref: unified namespace for metabolites and biochemical reactions in the context of metabolic models. Nucleic Acids Research 49:D570–D574.

41. Orth JD, Conrad TM, Na J, Lerman JA, Nam H, Feist AM, Palsson BØ. 2011. A comprehensive genome-scale reconstruction of *Escherichia coli* metabolism—2011. Molecular Systems Biology 7:535.

42. Xavier JC, Patil KR, Rocha I. 2017. Integration of Biomass Formulations of Genome-Scale Metabolic Models with Experimental Data Reveals Universally Essential Cofactors in Prokaryotes. Metabolic Engineering 39:200–208.

